# Cell Lysate Microarray for Mapping the Network of Genetic Regulators for Histone Marks

**DOI:** 10.1101/230466

**Authors:** Li Cheng, Cheng-xi Liu, Shuang-ying Jiang, Sha Hou, Jin-guo Huang, Zi-qing Chen, Yang-yang Sun, Huan Qi, He-wei Jiang, Jing-fang Wang, Yi-ming Zhou, Daniel M Czajkowsky, Jun-biao Dai, Sheng-ce Tao

## Abstract

Protein, as the major executer for cell progresses and functions, its abundance and the level of post-translational modifications, are tightly monitored by regulators. Genetic perturbation could help us to understand the relationships between genes and protein functions. Herein, we developed a cell lysate microarray on kilo-conditions (CLICK) from 4,837 yeast knockout (YKO) strains and 322 temperature-sensitive mutant strains to explore the impact of the genome-wide interruption on certain protein. Taking histone marks as examples, a general workflow was established for the global identification of upstream regulators. Through a single CLICK array test, we obtained a series of regulators for H3K4me3 which covers most of the known regulators in *Saccharomyces cerevisiae*. We also noted that several group of proteins that are linked to negatively regulation of H3K4me3. Further, we discovered that Cab4p and Cab5p, two key enzymes of CoA biosynthesis, play central roles in histone acylation. Because of its general applicability, CLICK array could be easily adopted to rapid and global identification of upstream protein/enzyme(s) that regulate/modify the level of a protein or the posttranslational modification of a non-histone protein.

## INTRODUCTION

Genetic perturbation has been widely employed to illuminate gene function in biological research. With the increasing pace of technology advancement in genome editing approach and sequencing, several large-scale genetic perturbation approaches have been developed to map the genetic element regulators in cell phenotypes such as cell viability (Blomen, et al., 2015), cell differentiation (Jaitin, et al., 2016) and cell immune response (Parnas, et al., 2015), and *etc.* However, not all genetic perturbations resulted in obvious change of cell state. Protein, the major performer in cell processes, should be a better readout for genetic perturbation. However, genome-wide perturbation method to identify the network of genetic regulators on protein state, including its abundance and post-translational level, are still largely lacking. Recently, a method using random mutagenesis in haploid human cells is applied to protein measurements in individual cells (Brockmann, et al., 2017). However, this method relies on fluorescence activated cell sorter (FACS) and deep sequencing which require professional and complicate operation. Thus, there is a need to develop a quick and efficient method for globally probing of the genetic regulators and the underlying network on protein state.

Histones are one of the most important and conserved proteins in eukaryotic cells. They are subjected to many different types of post-translational modifications (PTMs), especially many novel PTMs discovered in the past few years (Huang, et al., 2015) including a wide range of lysine acylations, such as butyrylation (Chen, et al., 2007), crotonylation (Tan, et al., 2011) and β-hydroxybutyrylation (Xie, et al., 2016). Currently, there are more than 500 known histone marks (Huang, et al., 2015). Many of these histone marks play important roles in the regulation of DNA-related processes such as replication and transcription. However, how and by what proteins these histone marks are regulated remains largely unknown. The proteins affecting the dynamics of histone marks include “writers”, “erasers”, “readers”, and other regulators (Strahl and Allis, 2000). We define in this study all of these proteins as regulators of histone marks (RHMs), and classify them into two categories: positive-regulators or negative-regulators which perturbation could decrease or increase, respectively, the level of the corresponding histone mark.

There are several existing strategies for the discovery of RHMs. Some RHMs have been identified based on the chemical similarity of the particular modification. For example, since propionylation, butyrylation and crotonylation are structurally similar to acetylation, it was expected that the “writers” and “erasers” for acetylation (namely, histone acetyltransferases (HATs) and histone deacetylases (HDACs)), also regulate these other acylation states, and indeed, this was confirmed for crotonylation (Bao, et al., 2014; Sabari, et al., 2015). However, despite the hundreds of acylation modifications on histones, there are very few regulators have been identified to date. We anticipated that this number of regulators is too low to be responsible for the tight and pervasive regulation of diverse histone acylation states, and so expected that many novel regulators remain to be discovered.

An alternate strategy for RHM discovery is to screen thousands of random or targeted mutant cells such as an entire collection of a deletion library in an unbiased way by immunoblotting (Krogan, et al., 2003). Clearly however, it is not a trivial task to perform immunoblotting on thousands of samples. Recently, chromatin immunoprecipitation coupled with mass spectrometry (ChIP-MS) has been used to identify proteins bound at genomic regions associated with specific histones marks (Ji, et al., 2015). However, this method relies on stable interactions between proteins and histones, which may not be suitable to identify indirect regulators that do not bind the histones. To accelerate functional studies of histone marks, an efficient strategy for the fast and global identification of regulators at an affordable cost could be transformative.

*Saccharomyces cerevisiae* is a well-studied eukaryotic model organism. There are approximate 1,000 human disease related genes that are evolutionary conserved from *S. cerevisiae* to human (Heinicke, et al., 2007). Besides methylation and acetylation, a variety of other histone marks have also been identified on histones in *S. cerevisiae*, such as 2-hydroxyisobutyrylation (Dai, et al., 2014), succinylation and malonylation (Xie, et al., 2012). *S. cerevisiae* is a powerful system for genetic and epigenetic studies with many available tools and resources, such as the yeast knockout (YKO) collections that carry precise start-to-stop deletions of ~6,000 open reading frames (Giaever and Nislow, 2014) and a collection of temperature-sensitive (TS) mutants spanning 497 different essential genes (Ben-Aroya et al., 2008). Previously, the YKO collection has been successfully applied for the screening of histone PTM regulators by western blotting (Krogan, et al., 2003) or dot blotting (Han, et al., 2007). However, these strategies were significantly labor-intensive and time-consuming, limiting their general application for the discovery of histone regulators.

Microarrays, *e.g.* protein microarrays, are powerful tools for systematic analysis/discovery (Zhu, et al., 2001). Notably, microarrays enable proteome-wide analysis while consuming tiny amount of samples in a short period of time. For example, the reverse phase protein array (RPPA) (Paweletz, et al., 2001) contains hundreds to thousands of lysates from individual samples, *e.g.* tumors. By applying a set of antibodies for a specific pathway, the abundance of specific components of this pathway could be monitored quantitatively across many samples in a single test (Chan, et al., 2004).

In this study, we applied the protein microarray technology to fabricate a <u>c</u>ell <u>l</u>ysate m<u>ic</u>roarray on <u>k</u>ilo-conditions (CLICK) with the haploid YKO collection and the TS mutant strains to prepare an array enabling the examination of ~5,000 single proteins as potential RHMs in a single experiment. Using antibodies specific for H3K4me3 and H3K36me3, we demonstrate that most of the known regulators for these two histone marks could indeed be discovered in a single round of the CLICK assay. Next, using antibodies against many specific acyl modifications, we found that Cab4p and Cab5p, the final two enzymes of CoA biosynthesis, play central roles in acylation on histones. Since the CoA biosynthesis pathway is highly conserved, these RHMs may function to modulate histone acylation within eukaryotes more broadly. We anticipate that the CLICK array technology described here will become a generally applicable strategy for the systematical identification of RHMs and regulators of other proteins for both PTMs and abundance.

## RESULTS

### Overview and construction of the CLICK array

The schematic diagram of the CLICK array is depicted in **Figure 1A**. The collection containing thousands of certain gene deletion or mutant are cultured in certain condition. Different cell lysates represent different states for certain protein. Taking the histone mark as an example, in some cell lysates with positive-RHMs (such as gene B) or negative-RHMs (such as gene D) genetic perturbation, the histone mark level could be decreased or increased, respectively. Then these cell lysates are deposited onto a microarray which will be probed with two different primary antibodies simultaneously: one that targets histones generically and another that is specific for a particular histone mark. Subsequent probing of this array with fluorescently-labeled secondary antibodies specific for the primary antibodies enables a direct measurement of the levels of the particular histone mark and histone from the same spot. The ratio of the levels of histone mark/histone are calculated for each strain and compared to that of the WT. Variations such as batch-to-batch and array-to-array differences are greatly reduced with this dual color labeling strategy, thereby ensuring the reliability of the measurements. The positive-RHMs (the red spots on the merged array) and the negative-RHMs (the green spots on the merged array) are readily identified according to the rank of the ratio of histone mark/histone levels. With these regulators interaction information, we could easily draw a upstream regulation network for certain protein or protein PTMs.

**Figure 1.**
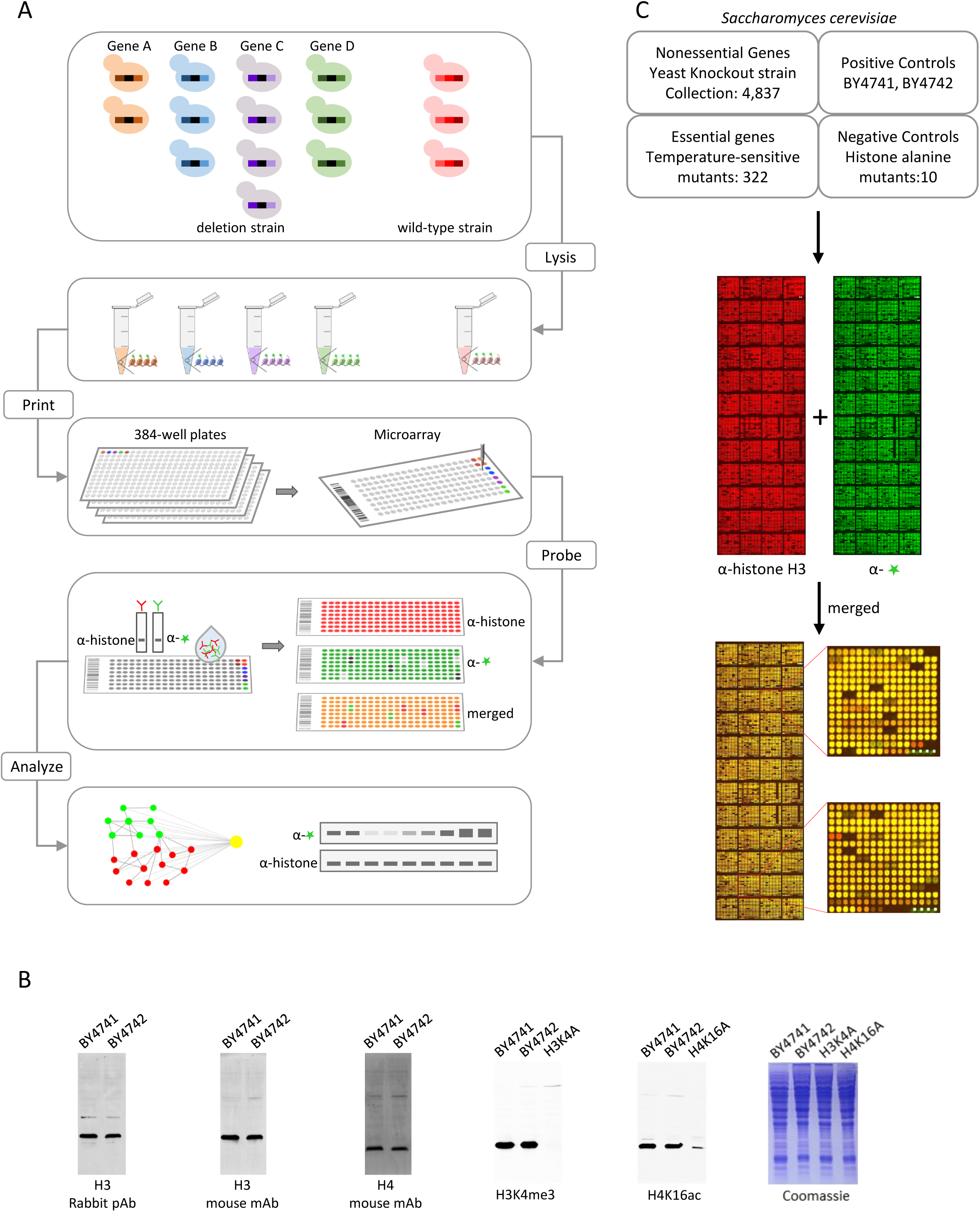
Schematic and Establishment of CLICK array. (A) Schematic illustration of CLICK array. The collection containing thousands of certain gene deletion or mutant are grow in certain condition. The orange, blue, purple and green shapes represent different genetic change strain each, and the red one is the wild-type strain. After collection and fractionation, the cell lysates are printed on microarray slides and following probe with specific antibodies. The strains with increased or decreased protein states can be recognized by the signal variation on microarray. At last, based on the microarray results and the previous knowledge, a upstream regulatory network could be constructed. (B) Determine antibodies that suitable for CLICK array. The criteria for selecting antibodies for CLICK array are as follows: only single band or near single band at expected size is allowed with total lysate by western blotting. Additionally, for histone mark-specific antibody, there should be no or very weak band in the corresponding alanine mutant strain. (C) The lysates of a total of 5,171 yeast cell lysates were spotted in duplication on a nitrocellulose coated glass slides (FAST™). After protein immobilization, the array was probe with an anti-histone H3 (Red), and a H3K4me3 specific antibody (Green).

One key factor of the CLICK array is the antibody specificity. Unlike assays such as western blotting that can tolerate some cross-reactivity in an antibody since the other proteins to which it binds can be separated from the protein of interest, this is not possible with the CLICK array: the complete collection of proteins of a strain are all immobilized on a single spot, and so any cross-reactions may lead to a strong collective signal. Therefore, preliminary assays to screen for CLICK-grade antibodies were first carried out. In particular, we examined the specificity of a given antibody by immunoblotting against the total lysates of WT strains (*S. cerevisiae* strains BY4741 and BY4742). A negative control consisting of a mutant histone protein in which the residue with the modification was mutated to alanine was included. That is, for example, H3K4A was included when testing an antibody for H3K4me3. An antibody was qualified for use in the CLICK array only if one major band at the expected size was present in the immunoblot, and at the same time no band or only very weak band was observed in the mutant sample. After screening a variety of modification-specific antibodies from different sources, we identified a set of antibodies that satisfied this criteria, including antibodies for H3K4me3, H4K16ac (**Figure 1B**), H3K36me3, H4K8ac, H3K9butyryl and H3S10p (**Figure S1A**). A similar examination of antibodies specific for histones H3 or H4 from a variety of manufactures, to be used as “loading controls” in the CLICK array, also identified several excellent candidates that only showed one major band in the total lysate from the WT strains (**Figure 1B**).

To assure the specificity of a histone mark-specific antibody, we next carried out dot blotting, which is more similar to the CLICK array since all of the cellular proteins are retained in one single spot. The ratios of the histone mark and histone (H3 or H4) were then measured. We found that among the six antibodies identified as highly specific in the western blotting assay, five showed similar levels of specificity in the dot blotting assay (**Figure S1B**): the antibody for H3K9butyryl was the sole candidate that failed in this assay (**Figure S1C**).

To construct the CLICK array, 4,873 nonessential deletion mutants from the haploid YKO collection and 322 temperature-sensitive essential genes mutants were included. As controls, we also included two WT strains, BY4741 and BY4742, and ten histone alanine mutants, namely H3K4A, H3K9A, H3S10A, H3K14A, H3K36A, H3K56A, H4K5A, H4K8A, H4K12A and H4K16A (**Figure. 1C**). The cell lysates were arrayed in duplicated spots onto a nitrocellulose-coated substrate slide.

To validate the quality of the resulting array, we probed it with a H3K4me3 specific rabbit polyclonal antibody and a H3 specific mouse monoclonal antibody, followed by Cy3- and Cy5-labeled secondary antibodies specific for rabbit and mouse, respectively (**Figure. 1C**). As expected, most of the spots displayed yellow, indicating that the ratios of H3K4me3 to H3 were constant across many mutant strains. That is, the majority of genes did not affect the level of H3K4me3 (**Figure. 1C**). Measuring the signal intensity of histone H3, the distributions of foreground and background signal intensity exhibited typical bell-shaped curves that were almost completely separate (**Figure S2**), indicating that the vast majority of the printed spots on the microarray contained substantial levels of histones.

### Identification of the network of RHMs of H3K4me3 by the CLICK array

To test the applicability of the CLICK array, we first examined for RHMs of H3K4me3, one of the best-characterized histone marks. The CLICK array was probed with the H3K4me3-specific antibody, and the fold-change of modification-to-histone level was determined as: for yeast strain X, the fold-change is (histone mark signal-X / histone signal-X) / (histone mark signal-WT/histone signal-WT), where the histone mark is H3K4me3 here, and WT is the BY4742 strain. The fold-changes for all strains on the CLICK array were then determined. Overall, we identified 21 positive-regulators and 61 negative-regulators (**Figure 2A**) and some of them were verified by western blot (**Figure 2B and 2C**). To illustrate the relationships among these RHMs, we combined our data with the known regulators prior to this study and analyzed their interactions using STRING (Szklarczyk, et al., 2017), where interacting proteins were grouped based on their GO classifications (**Figure 2D**). We anticipated to re-discovery previous known RHMs and some new RHMs. As expected, even though several known regulators were absent on the CLICK array, our data had covered all known complexes which were reported to regulate H3K4me3, including the COMPASS (*SWD3, BRE2, SWD1, SPP1* and *SDC1*) which mediates trimethylation of H3K4 (Dehe, et al., 2006), Paf1C (Polymerase Associated Factors 1 complex) (*CDC73*) which was shown to be required for H3K4me3, H3K36me3 and H2BK123ub (Krogan, et al., 2003; Wood, et al., 2003), the Rad6p-Bre1p-Lge1p complex (*RAD6* and *LGE1*) which is responsible for the ubiquitylation of H2BK123 (Hwang, et al., 2003), the BUR kinase complex (*SGV1*) (Laribee, et al., 2005) and CCR4/NOT complex (*POP2* and *MOT2*) (Laribee, et al., 2007) (**Figure 2D**). Surprisingly, we found that knockout of ARD1, encoding a subunit of protein N-terminal acetyltransferase NatA, which is known to regulate trimethylation of H3K79 (Takahashi et al.,2011), caused a 60% reduction of H3K4me3 (**Figure 2B**), implicating ARD1 as a previously unknown mediator of K79-K4 methylation crosstalk.

**Figure 2.**
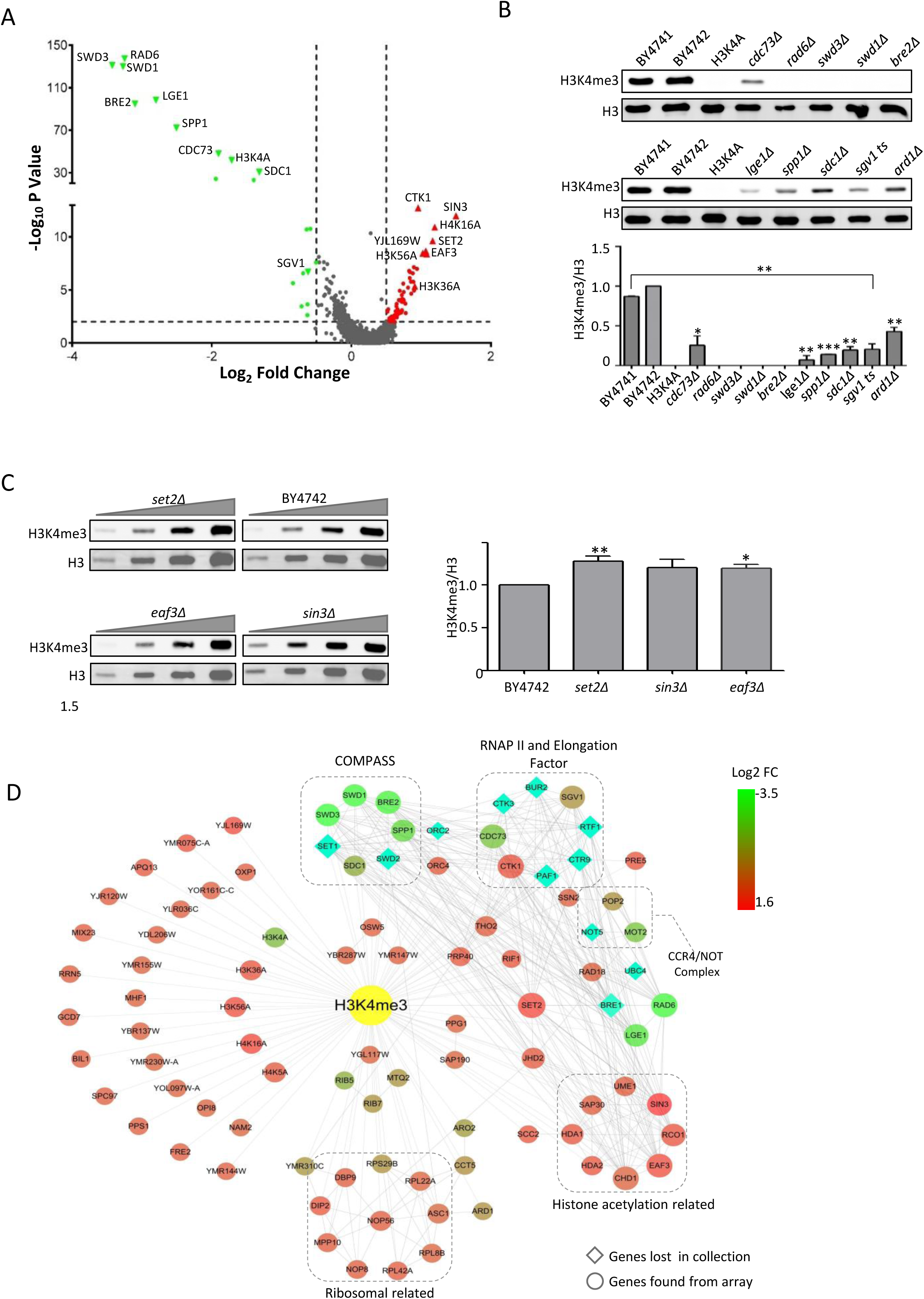
The Regulatory Network for H3K4me3 Resulting from CLICK array. (A) Volcano plot for H3K4me3. The green and red triangles indicate points-of-interest that display both large magnitude fold-changes (x axis) and high statistical significance (-log10 of p value, y axis). The dashed grey line of y axis denotes p = 0.01 and the dashed lines of x axis denote log2 of fold change =-0.5 or Log2 of fold change =0.5, respectively. This plot is colored such that those points having a fold-change less than 0.5 (log2 < 0.5) or above 0.5 and p value less than 0.01 simultaneously, are shown in green and red, respectively. (B) Validation of the positive-regulators by western blotting. Data represent mean fold change from three independent experiments ± SD. *P≤0.05, **P≤0.01, ***P≤0.001. (C) Validation of the negative-regulators. Serial dilutions of the total lysates were analyzed by western blotting. (D) The regulatory network for H3K4me3. 30 positive regulators and 63 negative regulators were further analyzed using STRING and visualized by Cytoscape.

We also identified several negative-regulators of H3K4me3 (**Figure 2C and 2D**), including the one previous reported, Ctk1p (Xiao, et al., 2007) and Jhd2p, a H3K4me3 demethylase (Liang, et al., 2007). Interestingly, the lysine to alanine mutant H3K36A and loss of Set2p, a histone methyltransferase of H3K36 (see below), both resulted in elevated H3K4me3, suggesting a previously unknown cross talk between these two histone marks. Moreover, four components of the Rpd3 histone deacetylase complex (*SIN3, EAF3, UME1, RCO1* and *SAP30*) and two HDA1 histone deacetylase complex members (*HDA1* and *HDA2*) showed negative regulation on H3K4me3 (**Figure 2D**). These observed regulations are consistent with the previous finding that histone deacetylase inhibitors could enhance the level of H3K4me3 (Nightingale, et al., 2007) and Gcn5 HAT complex enhances the level of H3K4me3 in transcribed coding sequences (Govind, et al., 2007). In addition, several proteins were enriched to ribosomal subunit components (*RPL22A, RPL42A, RPL8B, ASC1* and *RPS29B*) or those which are required for ribosomal subunit synthesis (*NOP56, NOP8, MPP10, DIP2, DBP9* and *YMR310C*) (**Figure 2D**). H3K4me3 is known to play a primary role in repression of ribosomal genes conjugating with H3S10p under multiple stress conditions (Weiner, et al., 2012). Thus, it is possible that H3K4me3 and certain ribosomal proteins may have a feedback regulation, which is worth further investigation.

Overall, through a single CLICK array experiment, a general view of RMHs for H3K4me3 could be obtained (**Figure 2D**). With their function annotations and previous studies, we can summarize the roles of known RHMs, and also explain the possible roles of the newly identified RHMs through their connections with the know RHMs.

### Identification of the network of RHMs of H3K36me3 by the CLICK array

To further validate the CLICK array, another histone mark, H3K36me3, was examined (**Figure S3**). Following a similar procedure to probe the array and analyze the data as described above, the fold-changes for all strains on the CLICK array were then determined. Overall, we identified 36 positive-RHMs and 54 negative-RHMs (**Figure S3A**) and some of them were verified by western blot (**Figure S3B**). Then we analyzed the interactions of these regulators as well as the known ones (**Figure S3C**).

H3K36me3 is associated with transcriptional elongation and is enriched throughout gene coding regions, peaking at the 3′ ends (Pokholok, et al., 2005). As expected, besides Set2p, the histone lysine methyltransferase (KMT) for H3K36me3, several regulators related to RNA polymerase (*RPB5* and *RPB7*) and transcription elongation factors (*CTK1, CTK3, CDC73, SGV1* and *TFG2*) were identified (**Figure S3C**). The result causing by deletion of YJL169W may due to its partial overlap with *SET2*. Many of these RHMs have been clarified for how to modulate H3K36me3 levels in previous studies (Wagner and Carpenter, 2012). For example, Set2p interacts with the phosphorylated C-terminal domain (CTD) of Rbp1p, the largest subunit of RNAPII, and this interaction is required for maintaining H3K36me levels (Kizer, et al., 2005). Two subunits of C-terminal domain kinase I (CTDK-I), Ctk1p and Ctk3p, are required for H3K36 methylation as their roles in the phosphorylation of CTD of Rbp1p (Youdell, et al., 2008). In addition, the absence of Paf1p, Ctr9p and Cdc73p, the three components of Paf1C, results in a loss of H3K36me3 and the deletion of *SGV1*, which is required for the proper recruitment of Paf1C was observed to have the same effect (Chu, et al., 2007). We also found several ribosomal related proteins negatively associated with the level of H3K36me3 (**Figure S3C**) but not H3K4me3. This may suggest that ribosomal proteins had different and specific roles in different histone marks. In fact, in mammal cells, the activation of KDM2A, demethylase for H3K36me1 and H3K36me2, could be inhibited when ribosome biogenesis is reduced under starvation (Tanaka, et al., 2010). Our results indicated that in yeast there may be a similar regulation yet to be discovered.

### Cab4p and Cab5p regulate the acetylation level of histone H4 on lysine 16

Given the capability of CLICK array to identify the regulators of well-studied modifications, we next apply the technology to discover modulators of H4K16ac, a functionally important but poorly studied histone mark.

Histone acetylation is known to be regulated by the opposing actions of HATs and HDACs. As the acetyl group has a negative charge, histone acetylation is expected to diminish the electrostatic affinity between histones and DNA, and thereby promote a more open chromatin structure that is more permissive to transcription (Shahbazian and Grunstein, 2007). H4K16ac is related to chromatin folding (Shogren-Knaak et al., 2006), heterochromatic silencing (Oppikofer, et al., 2011) and cellular lifespan (Dang, et al., 2009). In *S. cerevisiae*, H4K16 is acetylated primarily by the SAS complex, which consists of the catalytic subunit Sas2p and other two subunits, Sas4p and Sas5p (Suka, et al., 2002). H4K16 can also be acetylated by the essential HAT Esa1p, which is also responsible for acetylating other H4 tail lysines (Bird, et al., 2002). However, whether other factors are required for acetylation of H4K16 remains unknown.

A CLICK array was probed with specific antibodies against H4K16ac and H4 (**Figure 1B**) and the data analyzed following a similar process as described above, resulting in the identification of 188 positive-regulators and 26 negative-regulators (**Figures 3A**). Those with high fold-change were validation by western blotting (**Figure 3B**). To illustrate the relationships of these regulators, we analyzed their interactions and resulted some enriched sub-groups, including proteins related to aminoacyl-tRNA biosynthesis (*YDR341C, GLN4, YHR020W, GRS1, YNL247W* and *GUS1*), RNA polymerase subunits (*RPC11, RPC25, RPC37, RPB5* and *RPB8*) and proteins related to mRNA slicing (**Figure. 3C**). Interestingly, unlike H3K4me3 and H3K36me3, proteins related to ribosome showed positive regulations on H4K16ac (**Figure. 3C**). This suggested that even though these three histone marks were all related to transcriptional activity, they were obvious by different regulated.

**Figure 3.**
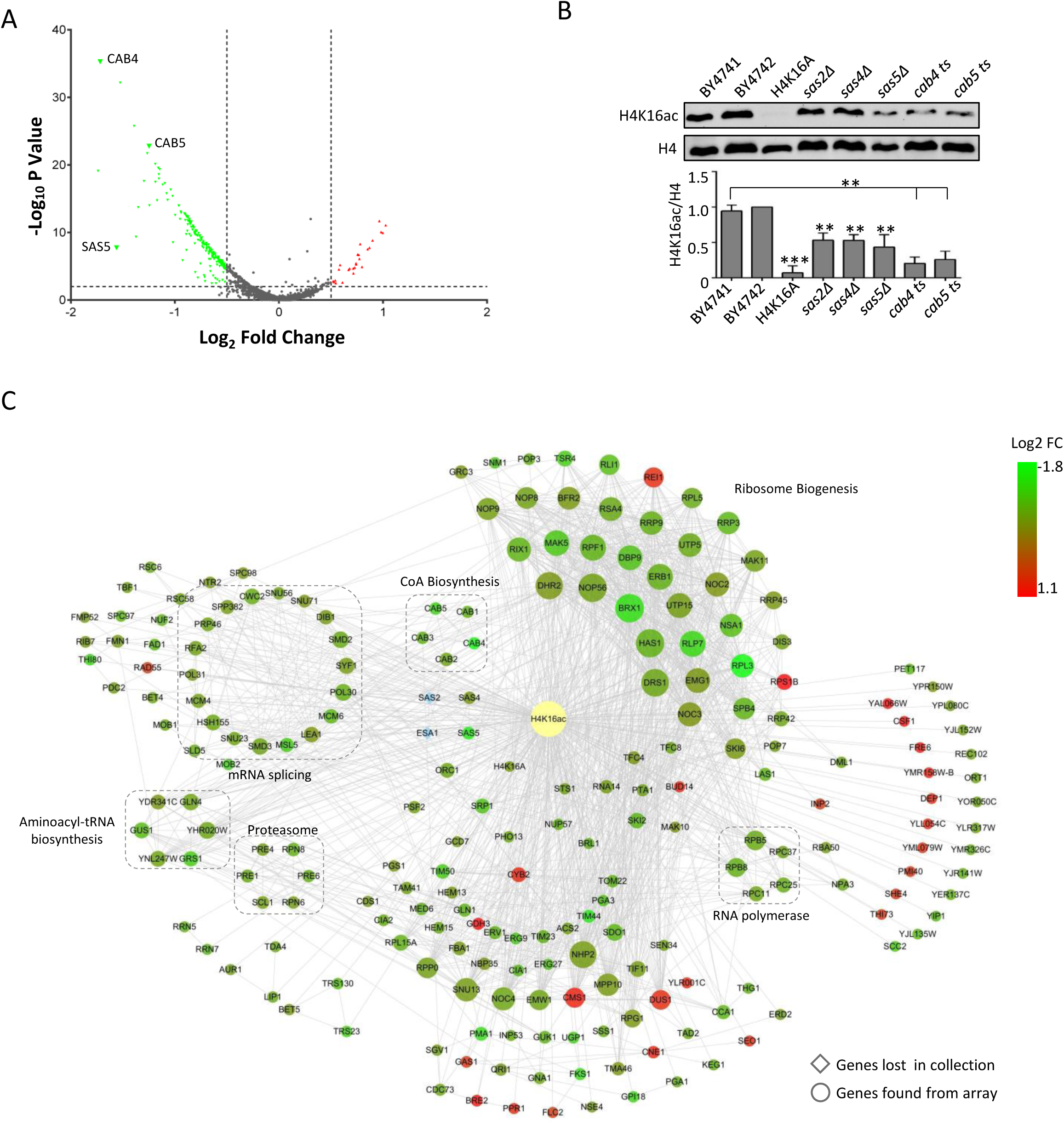
The Regulatory Network for H4K16ac Resulting from CLICK array. (A) Volcano plot for H3K4me3. The green triangles indicate points-of-interest that display both large magnitude fold-changes (x axis) and high statistical significance (-log10 of p value, y axis). (B) Validation of the positive-regulators by western blotting. Data represent mean fold change from three independent experiments ± SD. *P≤0.05, **P≤0.01, ***P≤0.001. (C) The regulatory network for H3K4me3. 191 positive regulators and 26 negative regulators were further analyzed using STRING and visualized by Cytoscape.

In addition, two essential genes, *CAB4* and *CAB5*, which together catalyze the last two steps of the coenzyme A (CoA) biosynthetic pathway, were on the top list of positive regulators (**Figure. 3C**). Immunoblot analysis suggested that Cab4p and Cab5p, in fact, had a more significant impact on H4K16ac levels than Cab1p, Cab2p and Cab3p, the other 3 genes in the CoA biosynthesis pathway consistent with the results of CLICK array (**Figure. 4A**).

**Figure 4.**
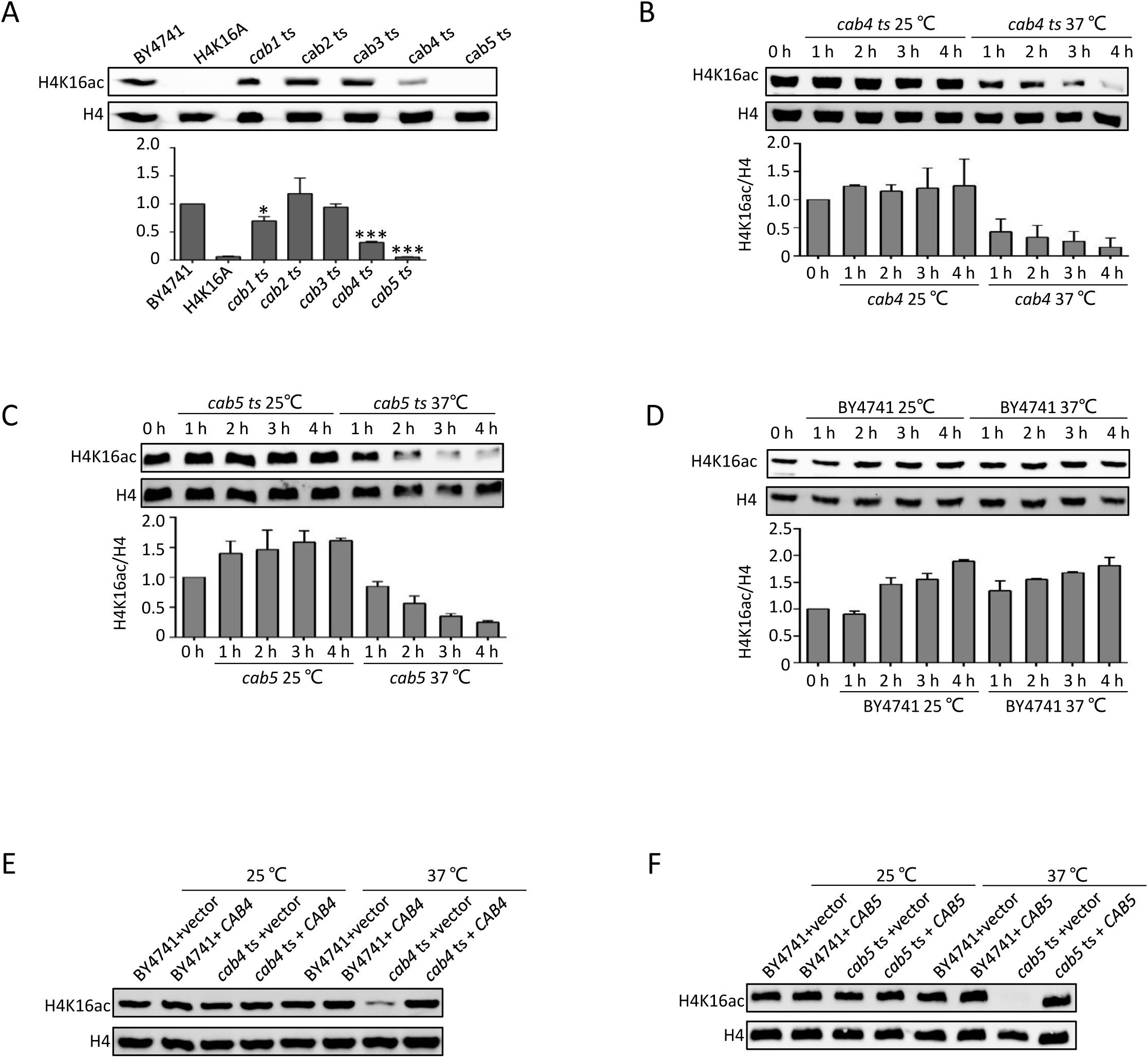
Cab4p and Cab5p are Up-regulators of H4K16ac. (A) Validation of the positive regulators taking part in the coenzyme A (CoA) biosynthetic pathway by western blotting. (B) -(D) H4K16ac level changes when shift from 25°C to 37°C. *Cab4* ts mutant (C), *cab5* ts mutant (D), and WT strain (E). The H4K16ac levels were measured at different time points. The cells were grow at 25°C to OD_600_ = 0.8, and then shifted to 25°C or 37°C for 1 h, 2 h, 3h and 4 h, and immunoblotted as indicated. (E)-(F) Wild type Cab4p and Cab5p could recover the H4K16ac level of *cab4* ts mutant (E), and the H4K16ac level of *cab5* ts mutant (F).

We then focused on Cab4p and Cab5p. To clearly demonstrate the effects of Cab4p and Cab5p on the H4K16ac modification, *cab4* and *cab5* ts mutants were shifted from the permissive temperature (25 °C) to the non-permissive temperature (37°C) in YPD medium for 4 h. The samples were collected throughout this 4-h period (at 1, 2, 3, and 4 h) and subjected to western blotting using the H4K16ac antibody. Clear loss of H4K16 acetylation was observed for both *cab4* and *cab5* ts mutants after the shift to 37°C for as short as 1 h (**Figures 4B, 4C**). Almost complete loss of the H4K16ac modification was observed after the mutants were shifted to 37°C for 4 h (**Figures 4B, 4C and 4D**). To further confirm the roles of Cab4p and Cab5p in regulating H4K16ac, WT *CAB4* and *CAB5* were complemented in *cab4* and *cab5* ts mutants, respectively. The growth defects of the *cab4* and *cab5* ts mutants at the non-permissive temperature were restored in these strains (**Figures S4A-D**). At the same time, the loss of H4K16ac observed in the original ts strains was also restored (**Figures 4E and 4F**). These results strongly suggest that Cab4 and Cab5 are two key regulators of H4K16ac.

### Cab4p and Cab5p play central roles in histone lysine acylation

Cab4p and Cab5p catalyze the last two steps of CoA biosynthesis (Olzhausen, et al., 2009): Cab4p adds the AMP moiety to 4′-phosphopantetheine forming dephospho-CoA, and then Cab5p phosphorylates the 3′-OH of the ribose to yield CoA. CoA biosynthesis is an essential pathway and highly conserved (Leonardi, et al., 2005). CoA is the precursor of acetyl-CoA and a variety of other acyl-CoA, such as crotonyl-CoA, propionyl-CoA and butyryl-CoA (**Figure 6A**). We postulated that Cab4p and Cab5p may not only play roles in H4K16 acetylation, but also be critical in the acetylation of other histone sites and, further, other types of acylations. To test this hypothesis, *cab4* and *cab5* ts mutants were shifted to 37 °C and samples were obtained at different time points. The cell lysates were analyzed by a variety of histone acylation antibodies, including H3K56ac, H4K8ac, H3K9bu, H3K14cr, H4K12cr, H4K12prop and H4K16bhb. In addition, two histone methylation specific antibodies H3K4me3 and H3K36me3 were also included. The amount of loaded proteins was determined by H3- or H4-specific antibodies. The results clearly showed that, similar to H4K16ac, there is a significant loss of all histone acylations tested but no change in the extent of histone methylations examined for both *cab4* (**Figure 5A**) and *cab5* ts mutants (**Figure 5B**), while no significant changes were observed in the WT strain BY4741 (**Figure 5C**). To further confirm the roles of Cab4p and Cab5p in regulating histone acylation, ts mutant strains complemented with a plasmid carrying either WT *CAB4* or *CAB5*, respectively, were also analyzed. As expected, the loss of the histone acylations were restored in strains at non-permissive temperature for both Cab4p (**Figure 5D**) and Cab5p (**Figure 5E**). We hypothesized that the loss of acylation may due to the depletion of CoA and acyl-CoAs in cab4 and cab5 ts mutants. In support of this argument, ultra-performance liquid chromatography-mass spectrometry (UPLC-MS) analysis of *S. cerevisiae* metabolites confirmed the significant loss of CoA, acetyl-CoA and butyryl-CoA in *cab4* and *cab5* ts mutants, and the loss could be at least partially restored by putting back the WT CAB4 and CAB5 (**Figure 6B-D**).

**Figure 5.**
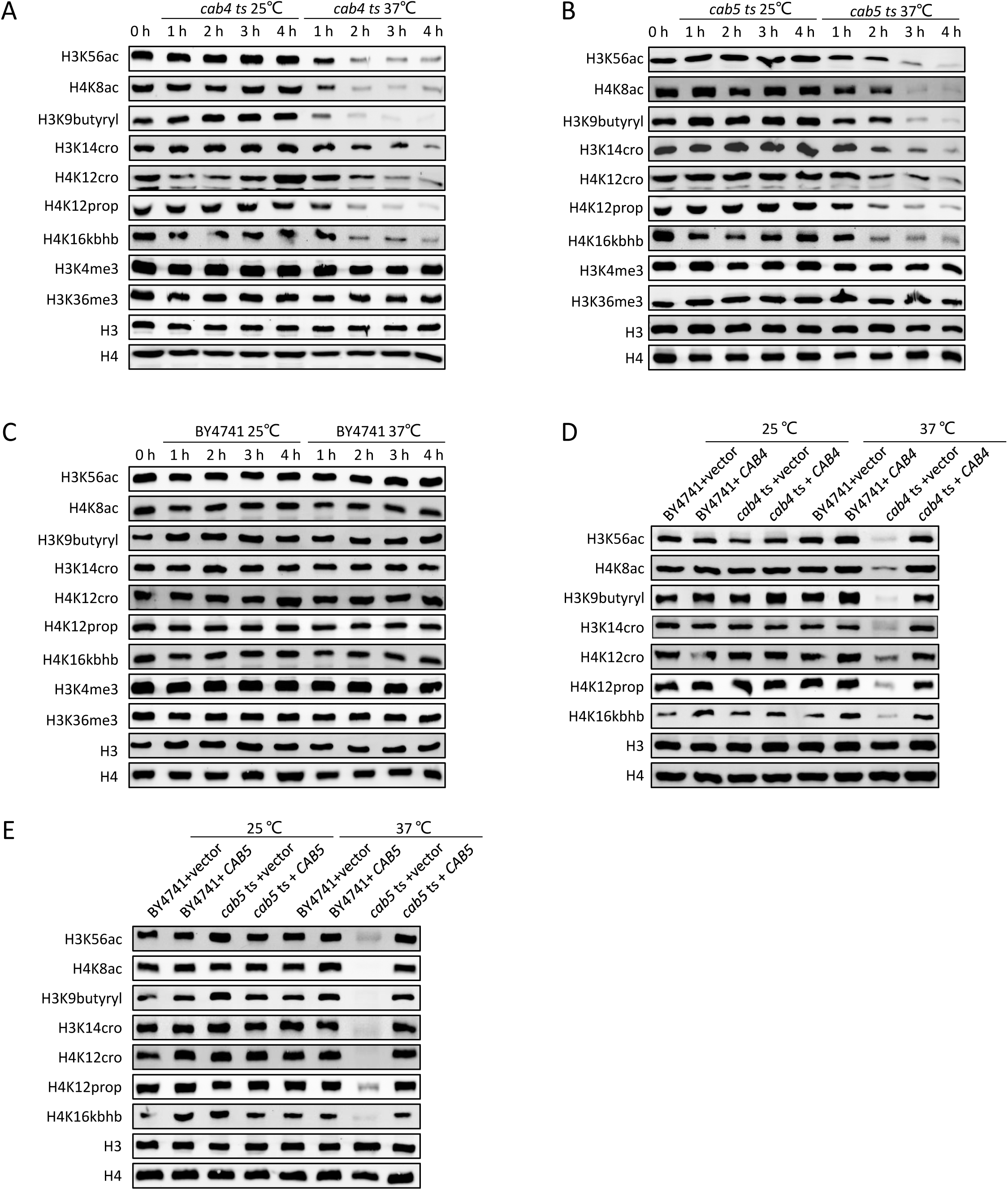
Cab4p and Cab5p Played Central Roles on Histone Lysine Acylation. (A)-(C) Inactivate Cab4p and Cab5p decreased the level of histone acetylation and other acylations in general. *Cab4* ts mutant (A), *cab5* ts mutant (B), and WT strain (C). (D)-(E) Wild type Cab4p and Cab5p could restore histone acylation in the ts mutants. *Cab4* ts mutant (D), and *cab5* ts mutant (E)

**Figure 6.**
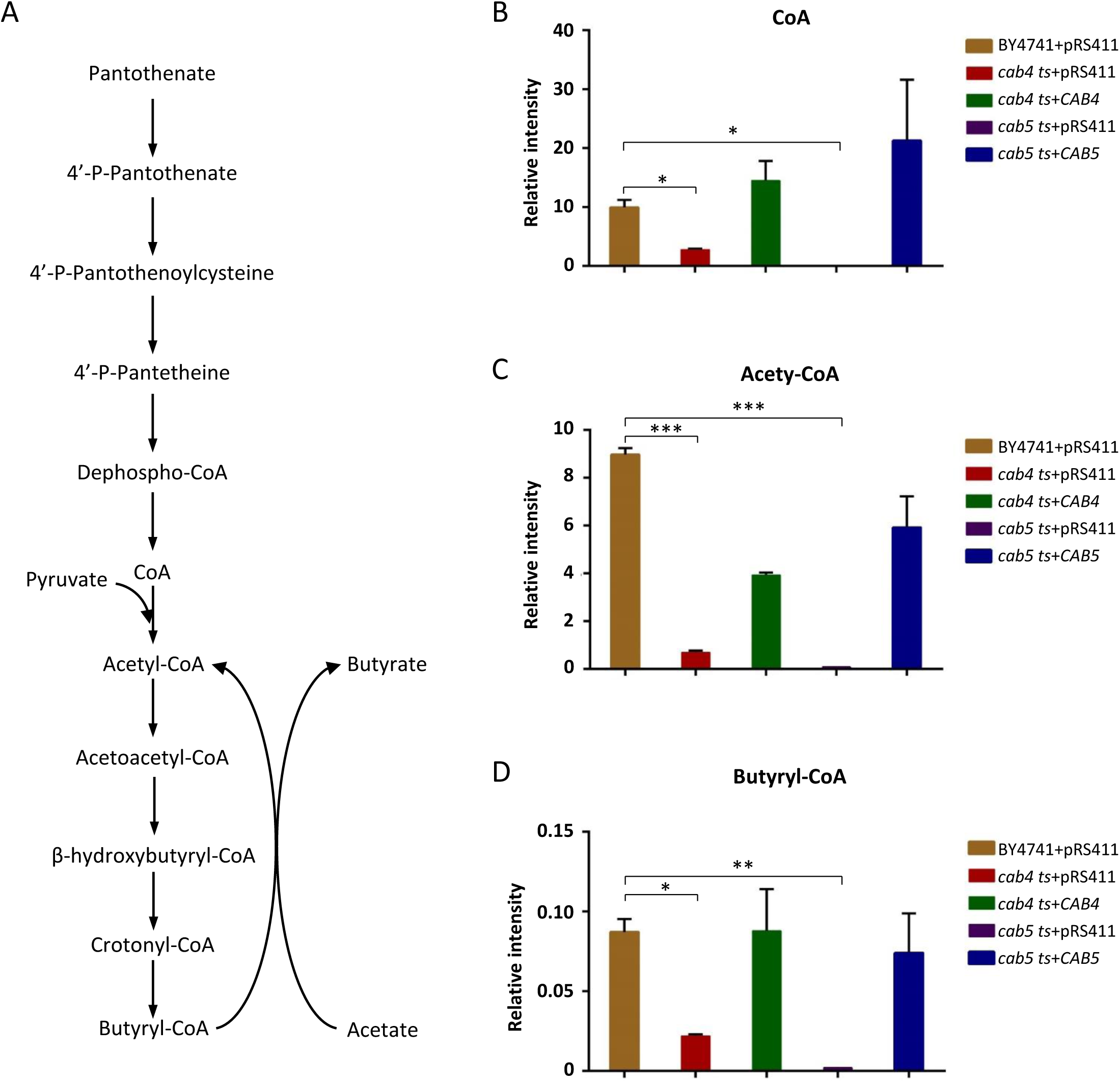
Cab4 and Cab5 Mutant lead the significant loss of CoA and Acyl-CoA. (A) The pathway of the biosynthesis of CoA and acyl-CoAs; (B)-(D) The intracellular level of CoA (B), acetyl-CoA (C) and butyryl-CoA (D) are maintained by Cab4p and Cab5p. Cells were grown at 25°C and then transferred to 37°C for 4 h. Metabolites were extracted and analyzed. The levels of CoA and acetyl-CoA were normalized by the amount of internal standard (Acetyl-1,2-^13^C_2_-CoA).

As *cab4* and *cab5* ts mutants affect the acylation of histone, we expected that *cab1, cab2* and *cab3* ts mutants would also affect histone acylation. Consistent with this, and supporting our results, previous work showed that reduced expression of Ppc1p, the homologue of Cab2p in *Schizosaccharomyces pombe*, caused diminished histone acetylation (Nakamura, et al., 2012). However, we only detected a modest decrease in the level of histone acylation in the *cab1* ts mutant but no in the *cab3* ts mutant (**Figures S5A**). Compared with *cab3* ts mutant, the levels of CoA and acetyl-CoA in *cab1* ts mutant showed significant loss, similar to those in *cab4* ts mutant (**Figures S5B and C**). For the *cab3* ts mutant, although it exhibited a growth defect at 37 °C (Figures S5D, F and G), we did not detect a statistically significant decrease in the level of CoA in a *cab3* ts mutant (**Figure S5F**), resulting undetectable loss of histone acylation. These results indicate that the influence on histone acylation modification of Cab4p and Cab5p not only result from their roles in CoA biosynthesis but also other unknown pathway.

### Cab4p and Cab5p response differently to stress

To determine whether *cab4* and *cab5* ts mutants have other influence on cellular function, we tested cell growth under different stress conditions. We found that both *cab4* and *cab5* ts mutants were sensitive to 200 mM hydroxyurea (HU) not only at 37°C but also at 25°C (**Figure S6A**). The inhibition of 200 mM HU could be restored for the *cab5* ts mutant, while not for the *cab4* ts mutant at the non-permissive temperature (**Figure S6A**). However, both *cab4* and *cab5* ts mutants were essentially insensitive to other DNA damage agents such as 10 μg/mL benomyl, 8 μg/mL camptothecin (CPT), 2 μg/mL nocodazole, 25 nM rapamycin and 3 mM H2O2 and to 100 J/m^2^ ultraviolet (UV) radiation (**Figure. S6B**). More interestingly, the *cab5* ts mutant exhibited a growth defect on YPG and YPGE medium at the permissive temperature but *cab4* ts mutant did not (**Figure. S6C**) and only the growth of *cab4* ts mutant could be restored on YPD containing 0.3% acetic acid at the non-permissive temperature (**Figure. S6D**). These results indicate that even though Cab4p and Cab5p were closely functioning in the same pathway, there are slight differences in metabolic regulations between them.

## DISCUSSION

In this study, we constructed a CLICK array with the cell lysates from *S. cerevisiae* strains of single gene knockouts or TS mutants. The array covers 82% of the open reading frames (ORFs) of *S. cerevisiae*. To realize the possibility for applying CLICK array for global discovery of genetic regulators for a given protein, we took histone marks as examples. By combining the array and histone mark-specific antibodies, we developed a general strategy enabling global identification of regulators of a given histone mark by a simple array probing. The networks of genetic regulators for H3K4me3, H3K36me3 and H4K16ac were identified through probing specific antibodies on the CLICK arrays. More importantly, using the CLICK array platform, we identified two key enzymes of CoA biosynthesis, Cab4p and Cab5p, that have a global impact on histone acylation.

### Advantages and disadvantages of CLICK array

As compared with existing strategies, such as Global Proteomic Analysis of *S. cerevisiae* (GPS) (Krogan, et al., 2003) and microwestern (Ciaccio, et al., 2010), the CLICK array has several advantages. Firstly, it enables proteome-wide identification of regulators for any protein or PTM, i.e., histone marks. Since the array covers 96% of the nonessential genes and 26% of the essential genes, we are able to screen 82% of the proteome of *S. cerevisiae* in a single experiment. Secondly, the screening process is fast, only requiring 14-15 h. In comparison, the western blotting based strategy, GPS, takes at least several months to screen the regulators of a single histone mark. Thirdly, the CLICK array is highly reliable. The dual-color modification/histone labeling strategy, similar to that commonly used in DNA microarrays (Churchill, 2002), reduces many of the variations encountered in protein microarray experiments, such as batch-to-batch, array-to-array, or spot-to-spot variations. Simply calculating the Cy3/Cy5 ratio for each spot also greatly simplifies the data analysis. Fourthly, the CLICK array based strategy is physiological relevant. To make the CLICK array, the whole cell lysate is denatured and printed onto the array directly. Theoretically, most, if not all, of the proteins from a strain will be immobilized on a single spot, providing a “snapshot” of the particular physiological proteomic status. We thus expect that the level and the pattern of the histone marks are well-retained on the CLICK array. Likewise, therefore, any regulators identified are anticipated to be physiologically important. Lastly, the CLICK array is a high-throughput strategy. Usually, it consumes 0.3−0.5 nL of sample per spot. Thus, the cell lysate prepared from one batch of 1 mL culture is enough to print 1,000 to 10,000 CLICK arrays, which lowers the cost for a single microarray to negligible. The printed arrays can be stored and later probed individually or in batch whenever necessary.

To make the CLICK array strategy generally applicable, several limitations have to be overcome. Firstly, it is not easy to obtain well-qualified antibodies, for example, a histone mark-specific antibody suitable for CLICK array assay. Even though there are more than 1,000 commercially available antibodies for various histone marks (Rothbart, et al., 2015), recent studies have revealed many unpredicted and alarming observations regarding the properties of the antibodies, including non-specificity, strong influence by adjacent PTMs, and inability to distinguish the modification state on a particular residue (*e.g.*, mono-, di-, or tri-methyl lysine) (Bock, et al., 2011; Egelhofer, et al., 2011). However, this is indeed now a well-recognized problem, with many focused efforts working on its resolution (Hattori, et al., 2013; Rothbart, et al., 2015). We anticipate that more histone mark-specific antibodies of CLICK array-quality will soon emerge. Secondly, the lysates on the current CLICK array represent only a single culture condition. Since histone marks can change under different culture conditions, some modifications may be absent or under represented on the current CLICK array. This possibility could be easily overcome by preparing several sets of cell lysates at different culture conditions, such as investigating mid-log vs stationary phase, adding sodium butyrate to improve the acetylation level (Davie, 2003), or adding nocodazole to improve the phosphorylation level (Baker, et al., 2010). All of these lysates could then be spotted onto a single array or a set of arrays, enabling a more profound screening of regulators for histone marks. Lastly, many of the regulators identified by the CLICK array could be indirect regulators, that is neither “writer” nor “eraser”. It may be difficult to further dissect their molecular mechanisms. To overcome this issue, bioinformatics, proteomics or other techniques, *e.g.* protein microarray could be applied to provide homolog or interacting proteins.

### Toward a comprehensive regulatory and functional network for histone marks

The data set presented here provides a comprehensive view of RHMs for several histone marks. For studying their functional roles in biological progress such as transcription, specific histone residue mutant could help us to know which genes they could influence. With powerful genomic technologies such as DNA microarray and RNA sequencing, it is easy to globally reveal the downstream consequence of an alteration on a given gene or protein. Thus, combining our CLICK array results and gene expression dataset, we could, for the first time, obtain a network which contains both the upstream regulators determining the abundance or PTM of certain protein, i.e., histone mark and the downstream global consequence in response to the corresponding specific interruption, i.e., histone residue mutant, in another word, a “complete” regulation picture of a given protein or PTM (**Figure 7, S8 and S9**). For example, in H3K4A mutant strain (GSE29059) (Jung, et al., 2015), the down expression genes were clustered to ribosomal biogenesis while the up expression genes were majorly located in mitochondria and related to energy derivation. However, the expression levels of the upstream regulators of H3K4me3 that we identified have not been obviously altered, suggesting that the transcription of these upstream regulators are not affected by H3K4A mutant or be masked by backup mechanisms.

**Figure 7.**
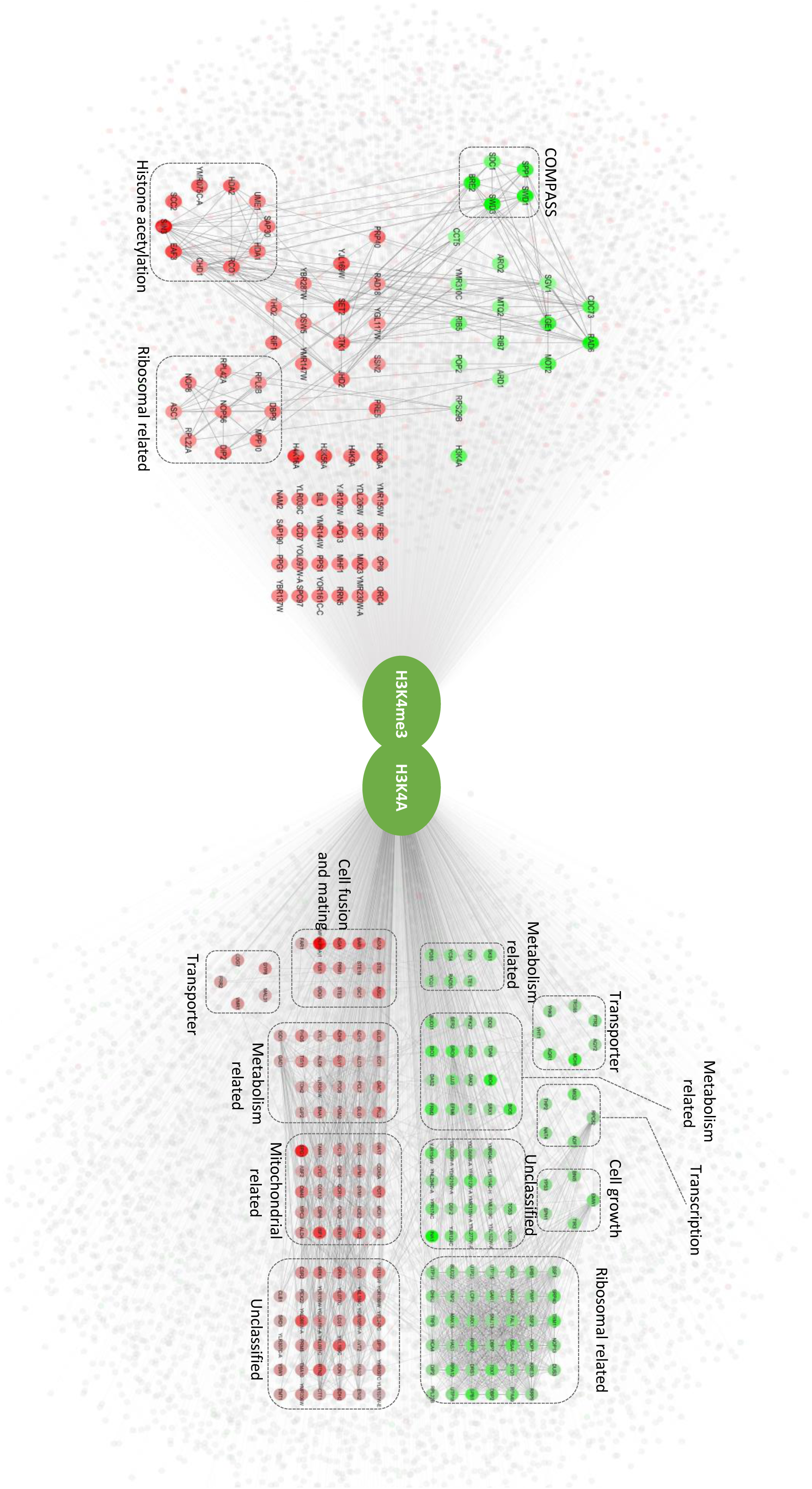
A complete network containing both upstream and downstream regulators for H3K4. The upstream regulators were collected through the combination of the CLICK array data and previous knowledge. The downstream results were collected through the expression data of H3K4A mutant, which is publically available.

### Crosstalk between H3K4me3 and the other two histone marks H3K36me3 and H4K16ac

In this study, we have built three global regulating networks with both the upstream and downstream regulators for H3K4me3, H3K36me3 and H4K16ac (**Figure 7, S8 and S9**). We found there are several regulators shared among these histone marks (**Figure S7**).

For H3K4me3 and H3K36me3, by CLICK array we found they shared 4 positive-regulators, 6 negative-regulators, and 4 regulators demonstrated the very reverse effect for these two histone marks (**Figure S7A**). Since some of the known regulators of H3K4me3 and H3K36me3 were missing from the CLICK array, these known regulators were then put together with the regulators that we identified on the CLICK array and subjected for cross talk analysis (**Figure S7B**). Indeed, Paf1C and BUR kinase complex has been reported to be required for H3K4me3 and H3K36me3 (Chu, et al., 2007; Krogan, et al., 2003; Laribee, et al., 2005). It is worth noting that 4 positive-regulators for H3K36me3 cause negative effect on H3K4me3. Previous study demonstrated that the deletion of *CTK1* elevated H3K4me3 level and this is related neither to COMPASS and Paf1C nor to Rad6/Bre1-mediated histone H2B monoubiquitination (Wood, et al., 2007). The interaction between Set2p and Pol II is due to phosphorylation of Pol II CTD by the CTK kinase (Xiao, et al., 2003). Our results indicates that the regulation of H3K36me3 by CTK1 results in its negative regulation role on H3K4me3. In addition, proteins associated with histone deacetylation, Sin3p and Eaf3p, were found to affect H3K4me3 levels (**Figure 2C**). Several studies have showed that H3K36me could be recognized by Eaf3p and recruits the Rpd3S deacetylase complex, and leads to deacetylation of H3 (Carrozza, et al., 2005; Joshi and Struhl, 2005; Keogh, et al., 2005). H3K4 methylation also facilitates histone acetylation (Wang, et al., 2009) suggesting a positive feedback loop between histone acetylation and H3K4 trimethylation. Both H3K4 methylation and H3K36 methylation are considered to be associated with transcriptionally active genes and H3K4me3 peaks at the beginning of the transcribed portions of genes while H3K36me3 is abundant throughout the coding region, peaking near the 3′ ends (Pokholok, et al., 2005). Recruitment of deacetylase by Set2p mediates H3K36 methylation, prevents spurious transcription within coding regions (Lee and Shilatifard, 2007), and may eventually lead to the loss of H3K4me3 to inhibit inappropriate initiation within coding regions. Altogether, our results and previous studies suggest a strong crosstalk between H3K4me3 and H3K36me3 through their different roles in histone acetylation regulation.

For H3K4me3 and H4K16ac, we found they shared 3 positive-regulators, and 10 regulators demonstrated the very reverse effect for these two histone marks (**Figure S7C and D**). In fission yeast, loss of Leo1p, a component of Paf1C, would lead to reduced level of H4K16ac at heterochromatin boundary regions (Verrier, et al., 2015). The recruitment of Paf1C is depend on the phosphorylation of the C-terminal repeats (CTRs) of Spt5 by Sgv1p (Liu, et al., 2009). As Cdc73p is a component of Paf1C, we speculate that the decrease of H4K16ac level caused by deletion of *CDC73* and *SGV1* may due to the similar mechanism. Moreover, we also found in H4K16A mutant strain, the level of H3K4me3 had a modest increase, as well as in H3K56A and H4K5A mutant strains, but not for H3K9A, H4K8A and H4K12A, even though these residues could be actylated. Even though H3K4me3 can trigger acetylation and deacetylation of histone H3 and H4 in certain contexts (Latham and Dent, 2007) and several positive regulatory residues on histones are required for the implementation of normal levels of H3K4me3 (Nakanishi, et al., 2008), it is little known about the negative regulatory residues for H3K4me3. The CLICK array provides a method for systematically profiling and comparing of regulatory residues on histones for certain histone mark since most of the histone lysine to alanine mutants are avaliable (Dai, et al., 2008).

In addition, we found that the N-terminal acetyltransferase subunit Ard1p also affects H3K4me3 (**Figures 2B**). Ard1p is required for maintaining levels of H3K79me3 and H2B monoubiquitination (Takahashi, et al., 2011). As histone H3K4 and H3K79 methylation is dependent on H2BK123 monoubiquitination (Nakanishi, et al., 2009), Ard1p may similarly affect the trimethylation of H3K4 through regulation of H2BK123 monoubiquitination. The relationship between Ard1p and H3K4me3 needs to be further characterized.

### The roles of Cab4p and Cab5p in histone acylation

In conventional analysis, identification of new regulator is usually based on protein-protein interactions that may miss indirect ones (Ji, et al., 2015). In this study, we found two indirect regulators, Cab4p and Cab5p, have a significant impact on H4K16ac (**Figure 4**). They are responsible for catalyzing the last two steps of CoA biosynthesis (Olzhausen, et al., 2009). CoA is an essential cofactor for a large number of enzymes involved in the transfer of acyl groups and the metabolism of carboxylic acids and lipids in all organisms (Leonardi, et al., 2005). Pantothenate is the donor for CoA biosynthesis. Cab4p adds the AMP moiety to 4′-phosphopantetheine forming dephospho-CoA, and then Cab5p phosphorylates the 3′-OH of the ribose to yield CoA. Acetyl-CoA, as the predominant CoA thioester, has been reported to have an influence on histone acetylation, and acetyl-CoA synthetase is also required for histone acetylation (Cai, et al., 2011; Takahashi, et al., 2006). As the depletion of CoA will have a global effect on the concentration of the acetyl-CoA, we hypothesized that the same effect on other histone acetylation sites, and even other kind of acyl modifications, might be observed. Consistent with this, we found that Cab4p and Cab5p affects all of the histone acetylation and acylation reactions that we have investigated, namely H3K56ac and H4K8ac, as well as butyrylation, propionylation, crotonylation, and β-hydroxybutyrylation. As expected, we found no effect on histone methylation in the *cab4* and *cab5* ts mutants (**Figure 5**), consistent with the idea that these mutants have their effect via a deficit of CoA. The UPLC-MS based metabolic analysis further supports our hypothesis that the reduced CoA level, resulting from the incomplete biosynthetic pathway, results in the depletion of acyl-CoA, which are the donors of histone acyl modifications. Therefore, the biosynthesis of CoA plays a critical role in histone acylation.

We note that the CoA biosynthesis pathway is conserved from bacteria to mammalian (Aghajanian and Worrall, 2002; Martinez, et al., 2014). In fact, the null mutants of enzymes in this pathway in *S. cerevisiae* can be complemented by their bacterial counterparts (Olzhausen, et al., 2009). CoA and acetyl-CoA participate in many biology processes and virtually every enzyme involved in glycolysis, fatty acid metabolism and the TCA and urea cycles is acetylated (Theodoulou, et al., 2014). However, we did not observe obvious differences in acetylation levels of non-histone proteins in *cab4* and *cab5* ts mutants as compared to that of WT strains (**Figure S10A**). The same result was observed when knocking out *coaE* in *E. coli*, which does not have histone proteins (**Figure S10B**). These results indicate that the depletion of CoA caused by the *cab4* and *cab5* ts mutations only significantly affects acylation of histones but not other non-histone proteins.

### Perspectives of the CLICK array

The CLICK array can be easily extended for wider applications. In this study, we investigated thousands of gene knockouts or TS mutants. Thousands of other conditions or cell states could be similarly investigated, such as chemical/drug treatment, culture conditions, or overexpression of single genes. Further, the application of the CLICK array is not limited to the discovery of regulators for histone marks, and it could be easily adopted for studying upstream regulators of other proteins or biological molecules, as long as high-affinity reagent is available. Though *S. cerevisiae* is a highly effective model for the CLICK array, this array is also readily applicable to other species, including human. To prepare a collection of cell lines of kilo-conditions, CRISPR/Cas9 (Shalem, et al., 2014), or RNAi (Echeverri and Perrimon, 2006) are two practical options. In addition, the histone marks and their specific antibodies are much more extensive in human than in *S. cerevisiae*. We anticipate that once the human CLICK array is available, it will be a very powerful tool for wide range of studies.

Taken together, to accelerate the discovery of regulators and the functional study for histone marks, taking advantage of the high-density array format and the specific antibodies, herein, we established the CLICK array strategy. Using *S. cerevisiae* as a model, we built regulatory networks for three histone marks and re-discover the majority of known regulators for H3K4me3 and H3K36me3. In a test for new factors, we showed that Cab4p and Cab5p, the last two enzymes in the CoA biosynthesis pathway play central and surprisingly specific roles in histone acylation through controlling the level of acyl-CoA. The CLICK array strategy is generally applicable for the identification of regulators of other histone marks and other proteins and it is also readily extensible for other species and biomolecules.

## AUTHOR CONTRIBUTIONS

S.-c.T. conceived and designed the project. J.-b.D. provided key reagents. L.C. and C.-x.L. performed antibodies screening. L.C., C.-x.L, S.H., Z.-q.C., J.-g.H. Y.-y.S. H.Q. and H.- w.J. constructed the microarray. L.C., J.-f.W. and Y.-m. Z. analyzed the data. S.-y.J. performed the complementation assay and phenotype assay. L.C., D.M.C., S.-c.T., and J.-b.D. wrote and revised the manuscript.

## ACKNOWLEDGMENTS

We thank Dr. Lei Feng of the Instrumental Analysis Center of Shanghai Jiao Tong University for her kind help with LC-MS analysis. We thank Jing Huang of Tsinghua University for his kind help with western blotting analysis. This study was supported in part by The National Key Research and Development Program of China Grant 2016YFA0500600, National Natural Science Foundation of China Grants 31670831 and 31370813.

## EXPERIMENTAL PROCEDURES

### Yeast strains

The yeast strains used for lysate microarray were taken from the yeast deletion collection (MATα haploid complete set) containing 4,837 nonessential deletion mutants (Winzeler, et al., 1999) and 322 temperature-sensitive mutants of essential genes (Ben-Aroya et al., 2008). At the same time, ten histone alanine mutants (H3K4A, H3K9A, H3S10A, H3K14A, H3K36A, H3K56A, H4K5A, H4K8A, H4K12A and H4K16A) from histone mutant library (Dai, et al., 2008) as negative controls.

### Antibodies

A complete list of antibodies used in our experiments were purchased from various sources (see **Supplementary Table 1**) or information on the antibodies. All antibodies were used at a dilution of 1:2,000 for western blotting, 1:200 for dot blotting and 1:1000 for microarray.

### Cell culture and lysate preparation

For gene deletion mutants, strains were inoculated from −80 °C stocks onto agar plates containing YPD + 200 μg/mL GENETICIN (Thermofisher), allowed to grow 48 h at 30 °C. Then inoculate yeast cells from agar plates to a 96-well 2 ml box in which every well contained 500 μL YPD liquid media and a 2 mm diameter glass ball, which facilitated the uniform growth. After 24 h of growth at 30 °C, transfer to 6 ml fresh YPD. Grown at 30 °C with vigorous shaking about 16 h, the culture should reach O.D._600_ 1.0-1.2. The cells were harvested by spinning at 3000 g for 5 min, and the cell pellets were washed with 500 μL water one time. The washed semi-dry culture was immediately stored in −80 °C freezer. For ts mutants, cells were grown in URA-/dextrose liquid media and reached to O.D._600_ 0.5-0.6 at 25 °C, then shifted to the nonpermissive temperature for 2 h. Frozen cells were thawed at room temperature, resuspended in 1 mL 0.1 M NaOH and incubated at room temperature for 10 min. After centrifugation, 60 μL SDS sample buffer (0.6 M Tris pH 6.8, 0.5% SDS, 20% glycerol, 2% β-mercaptoethanol) was added and the samples were vortexed briefly before heating at 95 °C for 10 min. The supernatants of yeast cell lysate were used for microarray and western blotting and stored at −20 °C.

### Cell lysate microarray fabrication

We transferred lysates to wells in 384-well polypropylene plates (10 μl/well). We used a contact-printing robotic SmartArrayer 48 microarrayer (CapitalBio) fitted with solid spotting pins to spot lysates onto FAST slides (Schleicher & Schuell BioSciences). Slides coated with a single nitrocellulose pad and each cell lysate spotted twice. The resulting microarrays were stored at −20 °C prior to use.

### Probing antibodies on the cell lysate microarray

CLICK array were blocked for 1 hr at room temperature with shaking in blocking buffer (3% BSA in Tris-buffered saline solution containing 0.1% Tween 20 detergent [pH 7.4]). After blocking, arrays were probed with 3 ml site-specific histone mark antibody (PTM Lab) and at the same time, H3 or H4 antibody from different species would be added as its loading control. After incubating overnight at 4 °C with shaking, arrays were washed three times with shaking in 1xTBST and then probed with 3 ml Cy5-donkey-anti-rabbit antibody and Cy3-donkey-anti-mouse antibody (1 μg/ml in blocking buffer) for 1 h at room temperature. After washing three times in TBST, arrays were dried in a SlideWasher (CapitalBio) and then scanned with a GenePix 4200A microarray scanner (Molecular Devices). Data were analyzed with GenePix Pro 6.0 (Molecular Devices).

### Extraction of CoA and acyl-CoA

*Saccharomyces cerevisiae* cells were grown in the SC-Met medium to O.D._600_ 1.4-1.6, and 70 ml of the culture was used for the metabolite extraction. After washed with ddH2O, the harvested cells were suspended in 500 μL of cold 80% (v/v) methanol (-40 °C). After ultrasonic crashing for 5 min, the cells were frozen at −80 °C for 30 min and then put on ice to thaw the cell suspension. Freeze-thaw cycles were repeated five times in order to extract the cells. The methanolic extracts were separated from the cell debris by centrifugation at 16,000 rpm at 4 °C for 15 min. The supernatant were evaporated to dryness and dissolved in 50 μL of 40% methanol containing 1 μg/mL Acetyl-1,2-^13^C_2_-CoA as internal standard for quantification.

### Mass spectrometry analysis

The analysis of CoAs was performed on the UPLC-MS/MS system consisting of a Waters Acquity UPLC System and AB Sciex Triple Quad™ 5500 system. Separation was performed on an ACQUITY UPLC HSS T3 analytical column (100×2.1 mm, 1.7 μm particle size) with eluent A (water, 5 mM Ammonium acetate, pH 6.9) and eluent B (acetonitrile). The optimized gradient program used in this study resulted in a total run time of 7 min: 0 - 0.5 min, 1%B; 4 min, 55%B; 4.5 min, 100%B; 5.5 min, 100%B; 5.6 - 7 min, 1%B. For MS detection, an electrospray ionization source was used and the ion source parameters for the positive mode were set as follows: source temperature, 600 °C; curtain gas (CUR), 35 psi; ion source gas 1 (GAS 1), 55 psi; ion source gas 2 (GAS2), 55 psi; collision gas (CAD), 8 psi; ion spray voltage (IS), 5500V; entrance potential (EP), 10V; collision cell exit potential (CXP1), 10V.

### Array analysis

We used GenePix Pro 6.0 to determine the median pixel intensities for individual features and background pixels in both Cy3 and Cy5 channels. All the data was analyzed by limma package in R. Two chanel color signals were normalized using “loess” algorithm (Smyth and Speed, 2003). We normalized the background-subtracted Cy5 intensities (site-specific histone mark antibody) to the level of histone (Cy3) at each feature by taking the ratio of Cy5 to Cy3. The value was used for all subsequent analysis. We calculated fold change with respect to BY4742 for each positive spots. P values was calculated using z-score test with non-parameter statistical method. Briefly, to avoid the fold-change bias, the median value and the median absolute deviation (MAD) value were taken as robust mean value and robust standard deviation, respectively. The all fold-change values were transformed to z-scores by the following formula: z-score = (fold-change-median(fold-change))/MAD(fold-change). P values were then calculated based on z-scores by applying standard normal distribution with two-tailed alternative hypothesis.

### Data Resources

Raw data files of the CLICK arrays profiling with H3K4me3, H3K36me3 and H4K16ac specific antibodies have been deposited on http://www.protein--microarray.com/ as accession number: PMDE230.

## Reference

Aghajanian, S., and Worrall, D.M. (2002). Identification and characterization of the gene encoding the human phosphopantetheine adenylyltransferase and dephospho-CoA kinase bifunctional enzyme (CoA synthase). Biochemical Journal 365, 13–18.

Baker, S.P., Phillips, J., Anderson, S., Qiu, Q.F., Shabanowitz, J., Smith, M.M., Yates, J.R., Hunt, D.F., and Grant, P.A. (2010). Histone H3 Thr 45 phosphorylation is a replication-associated post-translational modification in S. cerevisiae. Nat Cell Biol 12, 294–U103.

Bao, X., Wang, Y., Li, X., Li, X.-M., Liu, Z., Yang, T., Wong, C.F., Zhang, J., Hao, Q., and Li, X.D. (2014). Identification of ‘erasers’ for lysine crotonylated histone marks using a chemical proteomics approach. Elife 3, e02999.

Ben-Aroya, S., Coombes, C., Kwok, T., O’Donnell, K.A., Boeke, J.D., and Hieter, P. (2008). Toward a comprehensive temperature-sensitive mutant repository of the essential genes of Saccharomyces cerevisiae. Molecular cell 30, 248–258.

Bird, A.W., Yu, D.Y., Praygrant, M.G., Qiu, Q., Harmon, K.E., Megee, P.C., Grant, P.A., Smith, M.M., and Christman, M.F. (2002). Acetylation of histone H4 by Esa1 is required for DNA double-strand break repair. Nature 419, 411–415.

Blomen, V.A., Majek, P., Jae, L.T., Bigenzahn, J.W., Nieuwenhuis, J., Staring, J., Sacco, R., Van Diemen, F.R., Olk, N., and Stukalov, A. (2015). Gene essentiality and synthetic lethality in haploid human cells. Science 350, 1092–1096.

Bock, I., Dhayalan, A., Kudithipudi, S., Brandt, O., Rathert, P., and Jeltsch, A. (2011). Detailed specificity analysis of antibodies binding to modified histone tails with peptide arrays. Epigenetics 6, 256–263.

Brockmann, M., Blomen, V.A., Nieuwenhuis, J., Stickel, E., Raaben, M., Bleijerveld, O.B., Altelaar, A.F.M., Jae, L.T., and Brummelkamp, T.R. (2017). Genetic wiring maps of single-cell protein states reveal an off-switch for GPCR signalling. Nature 546, 307–311.

Cai, L., Sutter, B.M., Li, B., and Tu, B.P. (2011). Acetyl-CoA induces cell growth and proliferation by promoting the acetylation of histones at growth genes. Molecular cell 42, 426–437.

Carrozza, M.J., Li, B., Florens, L., Suganuma, T., Swanson, S.K., Lee, K.K., Shia, W.J., Anderson, S., Yates, J., Washburn, M.P., et al. (2005). Histone H3 methylation by Set2 directs deacetylation of coding regions by Rpd3S to suppress spurious intragenic transcription. Cell 123, 581–592.

Chan, S.M., Ermann, J., Su, L., Fathman, C.G., and Utz, P.J. (2004). Protein microarrays for multiplex analysis of signal transduction pathways. Nature medicine 10, 1390–1396.

Chen, Y., Sprung, R., Tang, Y., Ball, H., Sangras, B., Kim, S.C., Falck, J.R., Peng, J., Gu, W., and Zhao, Y. (2007). Lysine propionylation and butyrylation are novel post-translational modifications in histones. Molecular & cellular proteomics 6, 812–819.

Chu, Y., Simic, R., Warner, M.H., Arndt, K.M., and Prelich, G. (2007). Regulation of histone modification and cryptic transcription by the Bur1 and Paf1 complexes. The EMBO journal 26, 4646–4656.

Churchill, G.A. (2002). Fundamentals of experimental design for cDNA microarrays. Nat Genet 32 Suppl, 490–495.

Ciaccio, M.F., Wagner, J.P., Chuu, C.P., Lauffenburger, D.A., and Jones, R.B. (2010). Systems analysis of EGF receptor signaling dynamics with microwestern arrays. Nat Methods 7, 148–155.

Dai, J., Hyland, E.M., Yuan, D.S., Huang, H., Bader, J.S., and Boeke, J.D. (2008). Probing nucleosome function: a highly versatile library of synthetic histone H3 and H4 mutants. Cell 134, 1066–1078.

Dai, L., Peng, C., Montellier, E., Lu, Z., Chen, Y., Ishii, H., Debernardi, A., Buchou, T., Rousseaux, S., Jin, F., et al. (2014). Lysine 2-hydroxyisobutyrylation is a widely distributed active histone mark. Nature chemical biology 10, 365–370.

Dang, W., Steffen, K.K., Perry, R., Dorsey, J.A., Johnson, F.B., Shilatifard, A., Kaeberlein, M., Kennedy, B.K., and Berger, S.L. (2009). Histone H4 lysine 16 acetylation regulates cellular lifespan. Nature 459, 802–807.

Davie, J.R. (2003). Inhibition of histone deacetylase activity by butyrate. The Journal of nutrition 133, 2485S–2493S.

Dehe, P.M., Dichtl, B., Schaft, D., Roguev, A., Pamblanco, M., Lebrun, R., Rodriguez-Gil, A., Mkandawire, M., Landsberg, K., Shevchenko, A., et al. (2006). Protein interactions within the Set1 complex and their roles in the regulation of histone 3 lysine 4 methylation. The Journal of biological chemistry 281, 35404–35412.

Echeverri, C.J., and Perrimon, N. (2006). High-throughput RNAi screening in cultured cells: a user’s guide. Nature reviews Genetics 7, 373–384.

Egelhofer, T.A., Minoda, A., Klugman, S., Lee, K., Kolasinska-Zwierz, P., Alekseyenko, A.A., Cheung, M.S., Day, D.S., Gadel, S., Gorchakov, A.A., et al. (2011). An assessment of histone-modification antibody quality. Nature structural & molecular biology 18, 91–93.

Giaever, G., and Nislow, C. (2014). The yeast deletion collection: a decade of functional genomics. Genetics 197, 451–465.

Govind, C.K., Zhang, F., Qiu, H., Hofmeyer, K., and Hinnebusch, A.G. (2007). Gcn5 promotes acetylation, eviction, and methylation of nucleosomes in transcribed coding regions. Molecular cell 25, 31–42.

Han, J., Zhou, H., Horazdovsky, B., Zhang, K., Xu, R.M., and Zhang, Z. (2007). Rtt109 acetylates histone H3 lysine 56 and functions in DNA replication. Science 315, 653–655.

Hattori, T., Taft, J.M., Swist, K.M., Luo, H., Witt, H., Slattery, M., Koide, A., Ruthenburg, A.J., Krajewski, K., Strahl, B.D., et al. (2013). Recombinant antibodies to histone post-translational modifications. Nat Methods 10, 992–995.

Heinicke, S., Livstone, M.S., Lu, C., Oughtred, R., Kang, F., Angiuoli, S.V., White, O., Botstein, D., and Dolinski, K. (2007). The Princeton Protein Orthology Database (P-POD): A Comparative Genomics Analysis Tool for Biologists. PLOS ONE 2.

Huang, H., Lin, S., Garcia, B.A., and Zhao, Y. (2015). Quantitative proteomic analysis of histone modifications. Chemical reviews 115, 2376–2418.

Hwang, W.W., Venkatasubrahmanyam, S., Ianculescu, A.G., Tong, A., Boone, C., and Madhani, H.D. (2003). A conserved RING finger protein required for histone H2B monoubiquitination and cell size control. Molecular cell 11, 261–266.

Jaitin, D.A., Weiner, A., Yofe, I., Lara-Astiaso, D., Keren-Shaul, H., David, E., Salame, T. M., Tanay, A., Van Oudenaarden, A., and Amit, I. (2016). Dissecting Immune Circuits by Linking CRISPR-Pooled Screens with Single-Cell RNA-Seq. Cell 167, 1883–1896 e1815.

Ji, X., Dadon, D.B., Abraham, B.J., Lee, T.I., Jaenisch, R., Bradner, J.E., and Young, R.A. (2015). Chromatin proteomic profiling reveals novel proteins associated with histone-marked genomic regions. Proceedings of the National Academy of Sciences of the United States of America 112, 3841–3846.

Joshi, A.A., and Struhl, K. (2005). Eaf3 chromodomain interaction with methylated H3-K36 links histone deacetylation to Pol II elongation. Molecular cell 20, 971–978.

Keogh, M.C., Kurdistani, S.K., Morris, S.A., Ahn, S.H., Podolny, V., Collins, S.R., Schuldiner, M., Chin, K., Punna, T., Thompson, N.J., et al. (2005). Cotranscriptional set2 methylation of histone H3 lysine 36 recruits a repressive Rpd3 complex. Cell 123, 593–605.

Kizer, K.O., Phatnani, H.P., Shibata, Y., Hall, H., Greenleaf, A.L., and Strahl, B.D. (2005). A novel domain in Set2 mediates RNA polymerase II interaction and couples histone H3K36 methylation with transcript elongation. Molecular and cellular biology 25, 3305–3316.

Krogan, N.J., Dover, J., Wood, A., Schneider, J., Heidt, J., Boateng, M.A., Dean, K., Ryan, O.W., Golshani, A., Johnston, M., et al. (2003). The Paf1 Complex Is Required for Histone H3 Methylation by COMPASS and Dot1p: Linking Transcriptional Elongation to Histone Methylation. Molecular cell 11, 721–729.

Laribee, R.N., Krogan, N.J., Xiao, T., Shibata, Y., Hughes, T.R., Greenblatt, J.F., and Strahl, B.D. (2005). BUR kinase selectively regulates H3 K4 trimethylation and H2B ubiquitylation through recruitment of the PAF elongation complex. Current biology: CB 15, 1487–1493.

Laribee, R.N., Shibata, Y., Mersman, D.P., Collins, S.R., Kemmeren, P., Roguev, A., Weissman, J.S., Briggs, S.D., Krogan, N.J., and Strahl, B.D. (2007). CCR4/NOT complex associates with the proteasome and regulates histone methylation. Proceedings of the National Academy of Sciences of the United States of America 104, 5836–5841.

Latham, J.A., and Dent, S.Y. (2007). Cross-regulation of histone modifications. Nature structural & molecular biology 14, 1017–1024.

Lee, J.-S., and Shilatifard, A. (2007). A site to remember: H3K36 methylation a mark for histone deacetylation. Mutation Research-Fundamental And Molecular Mechanisms Of Mutagenesis 618, 130–134

Leonardi, R., Zhang, Y.M., Rock, C.O., and Jackowski, S. (2005). Coenzyme A: back in action. Progress in lipid research 44, 125–153.

Liang, G., Klose, R.J., Gardner, K.E., and Zhang, Y. (2007). Yeast Jhd2p is a histone H3 Lys4 trimethyl demethylase. Nature structural & molecular biology 14, 243–245.

Liu, Y., Warfield, L., Zhang, C., Luo, J., Allen, J., Lang, W.H., Ranish, J., Shokat, K.M., and Hahn, S. (2009). Phosphorylation of the transcription elongation factor Spt5 by yeast Bur1 kinase stimulates recruitment of the PAF complex. Molecular and cellular biology 29, 4852–4863.

Martinez, D.L., Tsuchiya, Y., and Gout, I. (2014). Coenzyme A biosynthetic machinery in mammalian cells. Biochemical Society transactions 42, 1112–1117.

Nakamura, T., Pluskal, T., Nakaseko, Y., and Yanagida, M. (2012). Impaired coenzyme A synthesis in fission yeast causes defective mitosis, quiescence-exit failure, histone hypoacetylation and fragile DNA. Open biology 2, 120117.

Nakanishi, S., Lee, J.S., Gardner, K.E., Gardner, J.M., Takahashi, Y.H., Chandrasekharan, M.B., Sun, Z.W., Osley, M.A., Strahl, B.D., Jaspersen, S.L., et al. (2009). Histone H2BK123 monoubiquitination is the critical determinant for H3K4 and H3K79 trimethylation by COMPASS and Dot1. The Journal of cell biology 186, 371–377.

Nakanishi, S., Sanderson, B.W., Delventhal, K.M., Bradford, W.D., Staehling-Hampton, K., and Shilatifard, A. (2008). A comprehensive library of histone mutants identifies nucleosomal residues required for H3K4 methylation. Nature structural & molecular biology 15, 881–888.

Nightingale, K.P., Gendreizig, S., White, D.A., Bradbury, C., Hollfelder, F., and Turner, B.M. (2007). Cross-talk between histone modifications in response to histone deacetylase inhibitors: MLL4 links histone H3 acetylation and histone H3K4 methylation. The Journal of biological chemistry 282, 4408–4416.

Olzhausen, J., Schubbe, S., and Schuller, H.J. (2009). Genetic analysis of coenzyme A biosynthesis in the yeast Saccharomyces cerevisiae: identification of a conditional mutation in the pantothenate kinase gene CAB1. Current genetics 55, 163–173.

Oppikofer, M., Kueng, S., Martino, F., Soeroes, S., Hancock, S.M., Chin, J.W., Fischle, W., and Gasser, S.M. (2011). A dual role of H4K16 acetylation in the establishment of yeast silent chromatin. EMBO J 30, 2610–2621.

Parnas, O., Jovanovic, M., Eisenhaure, T.M., Herbst, R.H., Dixit, A., Ye, C.J., Przybylski, D., Platt, R.J., Tirosh, I., Sanjana, N.E., et al. (2015). A Genome-wide CRISPR Screen in Primary Immune Cells to Dissect Regulatory Networks. Cell 162, 675–686.

Paweletz, C.P., Charboneau, L., Bichsel, V.E., Simone, N.L., Chen, T., Gillespie, J.W., Emmert-Buck, M.R., Roth, M.J., Petricoin, E., and Liotta, L.A. (2001). Reverse phase protein microarrays which capture disease progression show activation of pro-survival pathways at the cancer invasion front. Oncogene 20, 1981–1989.

Pokholok, D.K., Harbison, C.T., Levine, S., Cole, M., Hannett, N.M., Lee, T.I., Bell, G.W., Walker, K., Rolfe, P.A., Herbolsheimer, E., et al. (2005). Genome-wide map of nucleosome acetylation and methylation in yeast. Cell 122, 517–527.

Rothbart, S.B., Dickson, B.M., Raab, J.R., Grzybowski, A.T., Krajewski, K., Guo, A.H., Shanle, E.K., Josefowicz, S.Z., Fuchs, S.M., Allis, C.D., et al. (2015). An Interactive Database for the Assessment of Histone Antibody Specificity. Molecular cell 59, 502–511.

Sabari, B.R., Tang, Z., Huang, H., Yong-Gonzalez, V., Molina, H., Kong, H.E., Dai, L., Shimada, M., Cross, J.R., Zhao, Y., et al. (2015). Intracellular Crotonyl-CoA Stimulates Transcription through p300-Catalyzed Histone Crotonylation. Molecular cell 58, 203–215.

Shahbazian, M.D., and Grunstein, M. (2007). Functions of site-specific histone acetylation and deacetylation. Annual review of biochemistry 76, 75–100.

Shalem, O., Sanjana, N.E., Hartenian, E., Shi, X., Scott, D.A., Mikkelsen, T.S., Heckl, D., Ebert, B.L., Root, D.E., and Doench, J.G. (2014). Genome-scale CRISPR-Cas9 knockout screening in human cells. Science 343, 84–87.

Shogren-Knaak, M., Ishii, H., Sun, J.-M., Pazin, M.J., Davie, J.R., and Peterson, C.L. (2006). Histone H4- K16 acetylation controls chromatin structure and protein interactions. Science 311, 844–847.

Smyth, G.K., and Speed, T. (2003). Normalization of cDNA Microarray Data. Methods 31, 265–273.

Strahl, B.D., and Allis, C.D. (2000). The language of covalent histone modifications. Nature 403, 41–45.

Suka, N., Luo, K., and Grunstein, M. (2002). Sir2p and Sas2p opposingly regulate acetylation of yeast histone H4 lysine16 and spreading of heterochromatin. Nat Genet 32, 378–383.

Szklarczyk, D., Morris, J.H., Cook, H., Kuhn, M., Wyder, S., Simonovic, M., Santos, A., Doncheva, N.T., Roth, A., Bork, P., et al. (2017). The STRING database in 2017: quality-controlled protein-protein association networks, made broadly accessible. Nucleic acids research 45, D362–D368.

Takahashi, H., McCaffery, J.M., Irizarry, R.A., and Boeke, J.D. (2006). Nucleocytosolic acetyl-coenzyme a synthetase is required for histone acetylation and global transcription. Molecular cell 23, 207–217.

Takahashi, Y.H., Schulze, J.M., Jackson, J., Hentrich, T., Seidel, C., Jaspersen, S.L., Kobor, M.S., and Shilatifard, A. (2011). Dot1 and histone H3K79 methylation in natural telomeric and HM silencing. Molecular cell 42, 118–126.

Tan, M.J., Luo, H., Lee, S., Jin, F.L., Yang, J.S., Montellier, E., Buchou, T., Cheng, Z.Y., Rousseaux, S., Rajagopal, N., et al. (2011). Identification of 67 Histone Marks and Histone Lysine Crotonylation as a New Type of Histone Modification. Cell 146, 1015–1027.

Tanaka, Y., Okamoto, K., Teye, K., Umata, T., Yamagiwa, N., Suto, Y., Zhang, Y., and Tsuneoka, M. (2010). JmjC enzyme KDM2A is a regulator of rRNA transcription in response to starvation. EMBO J 29, 1510–1522.

Theodoulou, F.L., Sibon, O.C., Jackowski, S., and Gout, I. (2014). Coenzyme A and its derivatives: renaissance of a textbook classic. Biochemical Society transactions 42, 1025–1032.

Verrier, L., Taglini, F., Barrales, R.R., Webb, S., Urano, T., Braun, S., and Bayne, E.H. (2015). Global regulation of heterochromatin spreading by Leo1. Open biology 5.

Wagner, E.J., and Carpenter, P.B. (2012). Understanding the language of Lys36 methylation at histone H3. Nat Rev Mol Cell Bio 13, 115–126.

Wang, Z., Zang, C., Cui, K., Schones, D.E., Barski, A., Peng, W., and Zhao, K. (2009). Genome-wide mapping of HATs and HDACs reveals distinct functions in active and inactive genes. Cell 138, 1019–1031.

Weiner, A., Chen, H.V., Liu, C.L., Rahat, A., Klien, A., Soares, L., Gudipati, M., Pfeffner, J., Regev, A., Buratowski, S., et al. (2012). Systematic dissection of roles for chromatin regulators in a yeast stress response. PLoS biology 10, e1001369.

Winzeler, E.A., Shoemaker, D.D., Astromoff, A., Liang, H., Anderson, K., Andre, B., Bangham, R., Benito, R., Boeke, J.D., and Bussey, H. (1999). Functional characterization of the S. cerevisiae genome by gene deletion and parallel analysis. Science 285, 901–906.

Wood, A., Schneider, J., Dover, J., Johnston, M., and Shilatifard, A. (2003). The Paf1 complex is essential for histone monoubiquitination by the Rad6-Bre1 complex, which signals for histone methylation by COMPASS and Dot1p. The Journal of biological chemistry 278, 34739–34742.

Wood, A., Shukla, A., Schneider, J., Lee, J.S., Stanton, J.D., Dzuiba, T., Swanson, S.K., Florens, L., Washburn, M.P., Wyrick, J., et al. (2007). Ctk complex-mediated regulation of histone methylation by COMPASS. Molecular and cellular biology 27, 709–720.

Xiao, T., Hall, H., Kizer, K.O., Shibata, Y., Hall, M.C., Borchers, C.H., and Strahl, B.D. (2003). Phosphorylation of RNA polymerase II CTD regulates H3 methylation in yeast. Genes & development 17, 654–663.

Xiao, T., Shibata, Y., Rao, B., Laribee, R.N., O’Rourke, R., Buck, M.J., Greenblatt, J.F., Krogan, N.J., Lieb, J.D., and Strahl, B.D. (2007). The RNA polymerase II kinase Ctk1 regulates positioning of a 5’ histone methylation boundary along genes. Molecular and cellular biology 27, 721–731.

Xie, Z., Dai J Fau - Dai, L., Dai L Fau - Tan, M., Tan M Fau - Cheng, Z., Cheng Z Fau - Wu, Y., Wu Y Fau - Boeke, J.D., Boeke Jd Fau - Zhao, Y., and Zhao, Y. (2012). Lysine succinylation and lysine malonylation in histones. Molecular & cellular proteomics 11, 100–107.

Xie, Z., Zhang, D., Chung, D., Tang, Z., Huang, H., Dai, L., Qi, S., Li, J., Colak, G., Chen, Y., et al. (2016). Metabolic Regulation of Gene Expression by Histone Lysine beta-Hydroxybutyrylation. Molecular cell 62, 194–206.

Youdell, M.L., Kizer, K.O., Kisseleva-Romanova, E., Fuchs, S.M., Duro, E., Strahl, B.D., and Mellor, J. (2008). Roles for Ctk1 and Spt6 in regulating the different methylation states of histone H3 lysine 36. Molecular and cellular biology 28, 4915–4926.

Zhu, H., Bilgin, M., Bangham, R., Hall, D., Casamayor, A., Bertone, P., Lan, N., Jansen, R., Bidlingmaier, S., and Houfek, T.D. (2001). Global analysis of protein activities using proteome chips. Science 293, 2101–2105.

